# Pathway Analysis Through Mutual Information

**DOI:** 10.1101/2022.06.30.495461

**Authors:** Gustavo S. Jeuken, Lukas Käll

## Abstract

Pathway analysis comes in many forms. Most are seeking to establish a connection between the activity of a certain biological pathway and a difference in phenotype, often relying on an upstream differential expression analysis to establish the difference between case and control. This process usually models this relationship using many assumptions, often of a linear nature, and may also involve statistical tests where the calculation of false discovery rates is not trivial.

Here, we propose a new method for pathway analysis, MIPath, that relies on information theoretical principles, and therefore is absent of a model for the nature of the association between pathway activity and phenotype, resulting on a very minimal set of assumptions. For this, we construct a different graph of samples for each pathway and score the association between the structure of this graph and any phenotype variable using Mutual Information, while adjusting for the effects of random chance in each score.

Our experiments show that this method produces robust and reproducible scores that successfully result in a high rank for target pathways on single cell datasets, outperforming established methods for pathway analysis on these same conditions.

## Introduction

When conducting pathway analysis, one seeks to quantify the association between an alterations in phenotype and the activity in a biological pathway. This is typically done as a way to get an overview of large gene lists that are generated from high throughput biological experiments.

The traditional methods to conduct such analysis have been to first score the association between gene expression and phenotype, and then check for enrichment among the highest scoring genes [1–4]. Such methods hence require some type of precomputed statistics as an input, commonly a differential expression result, to make the association between phenotypes and pathway activity, typically a result of a linear model [5, 6].

There are also topology-based pathway analysis methods that take a more mechanistic view of pathways. However, as they require complete annotation of, not just the participating analytes, but also the way they interact, they are often seen as hard to use [7]. A more recent alternative is to first quantify the activity of a pathway in each sample, and then score its association to the phenotype [8–12].

Most pathway analysis methods were originally developed for bulk data, however, at least some are found robust also for single cell data [13]. Relatively recently other methods, such as iDEA [14] are developed directly with single cell data in mind.

What are the current limitations of the analysis of pathway analysis when analyzing single-cell data? Apart from the pathway databases lacking coverage and accuracy, most methods for pathway analysis test for differences between samples on case and control groups. This is an important limitation when it comes to single cell data, as samples are often taken from single individuals, and thus cannot be easily separated into case and control sets [15].

Here we describe MIPath, a pathway analysis method that defines a pathway specific random variable over the samples, and tests its association to any phenotypical classification over the samples. We will use concepts from information theory that allows us to score this relationship without making assumptions of linearity of the regulation of pathways.

## Methods

Our method MIPath consists of several interlinked steps. We 1) construct a Nearest Neighbor graph from the expression values in a pathway, then 2) we use those graphs to separate the samples into modules using a module detection algorithm. Subsequently, 3) the adjusted mutual information between the phenotype and the modules is calculated. Each of these steps are described in detail in the subsections below.

### Nearest Neighbors Graph

A *k* Nearest Neighbor (NN) graph can be constructed from any multivariate data by connecting elements *i* and *j* with a directed edge if *j* is one of the *k* points with the smallest distance to *i*. Constructing such a graph, can be computationally expensive if a large number of data points are involved, but the estimation of the approximate *k* nearest neighbors can be done much more efficiently [16].

Once the graph is constructed, we can further add informative features such as edge weights. A similarity measure based on the number of shared nearest neighbors (SNN) has been shown to perform better than traditional similarity measures in high dimensional spaces [17] and this approach has already been shown to work well for gene expression [18]. Conveniently, the number of shared neighbors between nodes can be efficiently calculated using a known property of adjacency matrices [19, p229]. If *A* is the adjacency matrix of the directed NN graph, then *A*^*T*^ is the same graph with the direction of edges reversed, and so *B* = *A* · *A*^*T*^ gives us an adjacency matrix from where we can extract the SNN information, that is because *B*_*ij*_ is the number two-step paths nodes *i* and *j*, the first taken in *A* and the second in *A*^*T*^. This is the same as the number of shared neighbors of *i* and *j* in *A*. To convert the number of shared neighbors to a similarity metric, we simply need to divide this number by the total number of neighbors of *i*, which in this case is exactly *k*.

### Leiden algorithm for module detection

If the nodes of a graph are separated into groups, its modularity is defined as the difference between the fraction of edges that fall within the groups and the expected fraction of these that are expected with a random distribution of edges. More specifically, it can be measured as

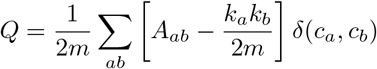

where *m* is the number of nodes, *A*_*ab*_ is an indicator variable of the existence of an edge between nodes *a* and *b, k*_*a*_ is the degree of node *a* and *δ*(*c*_*a*_, *c*_*b*_) is another indicator for the group assignment of nodes *a* and *b* (*c*_*a*_ and *c*_*b*_) being the same [20]. This can be adapted to deal with edge weights by changing *A*_*ab*_ to the weight of the edge from *a* to *b*, and to a directed network by changing *k*_*a*_ and *k*_*b*_ to 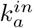 and 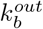, the in degree and out degree of nodes *a* and *b* respectively, and then multiplying the score by 2 [21].

The problem separating the nodes of a graph into modules in an optimal way can be viewed as an optimization problem on *Q*, with respect to the group assignments {*c*_*a*_, *c*_*b*_, …}. The solution to this problem is unfortunately not trivial. The Louvain method [22] first introduced a greedy approach to this problem that can be computed apparently in log-linear time [23]. The Leiden algorithm [24] improves on this method by providing further guarantees with a faster running time. The Leiden algorithm has also been used successfully by the Seurat package [25] on SNN networks, to find subpopulations of cells on single cell transcriptomics data.

### Adjusted Mutual information score

Mutual information (MI) is a measure of how much two random variables are associated to each other. More precisely, it measures the amount of information that is obtained about one of the variables when measuring the other. Importantly, it does not model the nature of the interaction between those variables, as it only looks at each variable’s entropy and their joint entropy. This makes MI an extremely useful measure of dependence where the shape of the interaction is not know, or can take multiple forms.

The MI of two random variables *X* and *Y* can be calculated as the Kullback–Leibler divergence (also known as relative entropy) between the joint distribution Pr(*X, Y*) and the product of the marginals Pr(*X*) Pr(*Y*):

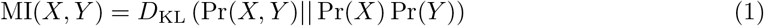

Another form of this expression is:

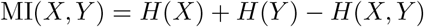

where *H*(*X*) is the entropy of a random variable *X*. This gives us a more intuitive interpretation of this measure. We can see that if the joint entropy is the same as the sum of the entropy of both variables, the MI is zero. Also, by using the chain rule, we can also see that the MI is maximal if the conditional entropies *H*(*X*|*Y*) or *H*(*Y* |*X*) are zero.

In our study, we will use MI to study the association between two partitions of a dataset, one that is based on gene expression, and another based on phenotype. As a metric, MI generally gives values in the range [0, +∞], which makes comparing these results difficult. However, it can be shown that the maximum MI between partitions *X* and *Y* is 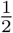 (*H*(*X*) + *H*(*Y*)) [26]. We can then define a Normalized Mutual Information (NMI) measure as:

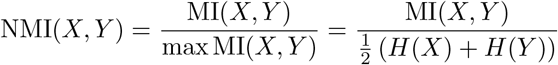

which is bounded to [0, 1], making comparisons easier. There is yet another problem in comparing NMI metrics, this time on the lower bound, because by random chance, MI will be higher when we have a bigger number of partitions [27]. We then need to define another measure, Adjusted Mutual Information (AMI), that takes into account this variation of the baseline:

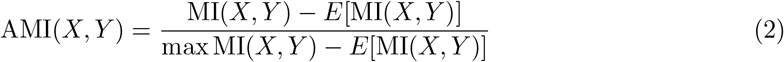

where *E*[MI(*X, Y*)] is the expected MI when both partitions *X* and *Y* are random. AMI will therefore be 1 if MI(*X, Y*) = max MI(*X, Y*) and 0 if MI(*X, Y*) = *E*[MI(*X, Y*)], and can also take very small negative values. This will be the measure used throughout this study.

Calculating *E*[MI(*X, Y*)] can be hard. However, here we have an advantage if using discrete classes, as by random the distribution of data on those classes will follow a generalized hypergeometric distribution. Vinh et al. [27] shows us how to calculate *E*[MI(*X, Y*)] under this model of randomness.

### Adjusted Conditional Mutual Information scoring

It is also possible to calculate a conditional mutual information MI(*X, Y* |*Z*), where *Z* is a third random variable. This is done simply by substituting the joint Pr(*X, Y*) by Pr(*X, Y* |*Z*) as well as the marginals Pr(*X*) and Pr(*Y*) by Pr(*X*|*Z*) and Pr(*Y* |*Z*), i.e., this can be calculated using the KL divergence as:

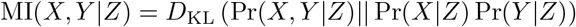

For a discrete variable 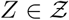, this can also be expressed as:

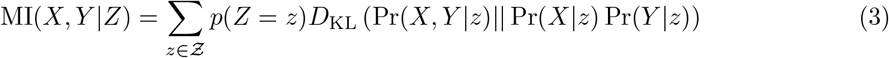

We can also define an AMI measure for the conditional probabilities:

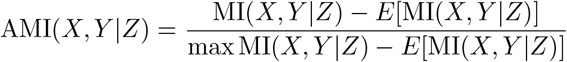

Where 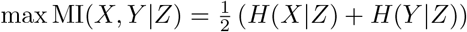 (following the above) and the calculation of *E*[*MI*(*X, Y* |*Z*)] can be found in the supplementary information.

To interpret the results of AMI(*X, Y* |*Z*), it is useful to compare it to AMI(*X, Y*). By looking at Equations 1 and 3 and the normalization that follows, we can see that situations where AMI(*X, Y* |*Z*) *<* AMI(*X, Y*) are those when for each value of *Z* the association of *X* and *Y* is weaker on average than AMI(*X, Y*), where we have no knowledge of the variable *Z*, and in this case *Z* can be viewed as a confounder. This difference between the mutual information and conditional mutual information is also called Interaction Information [28].

### Naïvely selected target pathways

Benchmarking the performance of a pathway analysis method is not a trivial task. For evident reasons, it is frequently done by interpreting the resulting pathway scores in the context of the experiment that was performed, as this is usually the circumstance on which these methods are required to perform. However important, this is a subjective measure of performance, thus prone to bias, where methods can be seen as successful both for presenting sound and expected results, or for elucidating new mechanisms.

Nguyen et al [29] argues for a reproducible measure for the success of pathway analysis methods. By selecting datasets with conditions that already have an associated pathway, e.g. Parkinson’s disease, one can benchmark methods by how they rank said pathway in their results. Bias can be avoided by selecting the datasets and target pathways before running any experiment. Here, we selected pathways based on the abstract of each set’s description. While this measure is very reductionist, it can be used across methods in a reliable and reproducible way, and thus is ideal for benchmarking of new methods. Note that we in several of the datasets made naïve assumptions about their nature. E.g. the major difference between metastases from melanoma in skin and liver in `E-CURD-84` might not be well described by the KEGG pathway “Melanoma”. However, here we stayed with our initial target.

### Datasets

Most datasets were downloaded from the Single Cell Expression Atlas [30]. Those datasets were selected according to the criteria on the section above, and their ascension number, associated publication, target pathway and tested phenotype can be seen on Table 1. The METABRIC dataset [31], was downloaded from European Genome-Phenome Archive under accession number EGAS00000000083.

**Table 1:**
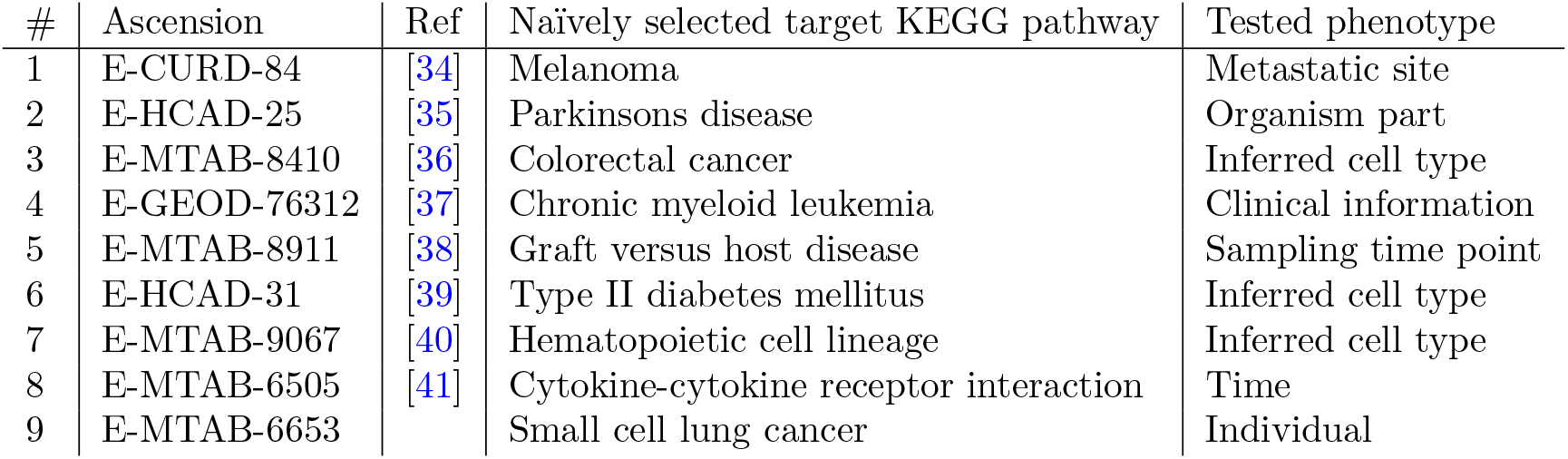
The Single cell Expression atlas datasets used in the comparison. Each dataset is listed with the phenotype variable we used for testing and the pathway we naïvely selected as a target pathway.

| # | Ascension | Ref | Naïvely selected target KEGG pathway | Tested phenotype |
| --- | --- | --- | --- | --- |
| 1 | E-CURD-84 | [34] | Melanoma | Metastatic site |
| 2 | E-HCAD-25 | [35] | Parkinsons disease | Organism part |
| 3 | E-MTAB-8410 | [36] | Colorectal cancer | Inferred cell type |
| 4 | E-GEOD-76312 | [37] | Chronic myeloid leukemia | Clinical information |
| 5 | E-MTAB-8911 | [38] | Graft versus host disease | Sampling time point |
| 6 | E-HCAD-31 | [39] | Type II diabetes mellitus | Inferred cell type |
| 7 | E-MTAB-9067 | [40] | Hematopoietic cell lineage | Inferred cell type |
| 8 | E-MTAB-6505 | [41] | Cytokine-cytokine receptor interaction | Time |
| 9 | E-MTAB-6653 |  | Small cell lung cancer | Individual |

As a source of pathway annotation, we have used the KEGG [32] pathways present in MSigDB [1], and Reactome [33] pathways downloaded from the project’s website.

## Results

### A pathway analysis method based on mutual information

For each pathway, we start by calculating the *k* approximate nearest neighbors of each sample, following Dong et al. [16], and using a euclidean distance metric on the normalized genes. Here, the distance metric used involves only the genes present on the pathway of interest, thus capturing only the sample structure present in the subspace defined by this pathway. This results in a directed graph that is then weighted using the SNN similarity.

At this point, we have a collection of graphs, one for each pathway, that have the exact same number of nodes and edges, with the only difference being that each is wired to represent neighborhoods on different pathway subspaces. To these, we apply the Leiden algorithm to detect modules present, and the resulting module assignments of the samples is used as a representation of the sample state for the pathway. Since the number of modules is a result of the algorithm, a pathway may have any different number of states.

The third step produces the pathway score by calculating the Adjusted Mutual Information between this pathway state of each sample and any other sample specific variable, such as annotations of phenotypes. Here we also note that we are not limited to studying case and control, but can score any discrete variable. Figure 1 presents an overview of the method.

**Figure 1:**
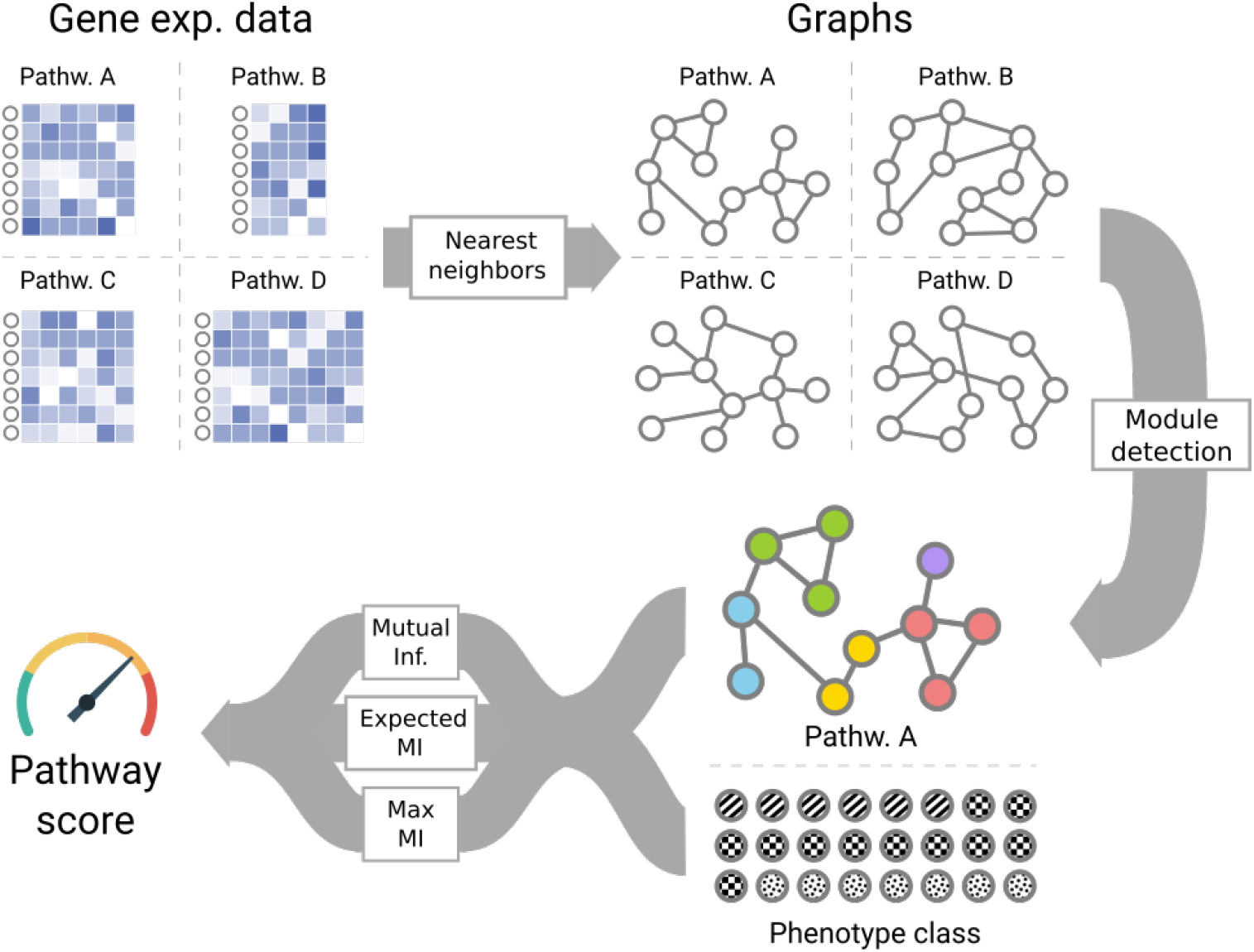
Method overview

Finally, we note that there is only one important parameter in the method, *k* number of neighbors, that presents a trade-off between local and global information, however the importance of this parameter is diminished by the use of the SNN similarity [17], as seen in the next section.

### Effects of parameter *k*

As stated, *k* is the only parameter of the model. The Approximate Nearest Neighbor algorithm used has an empirical complexity of *O*(*n*^1.14^) on the number of samples [16], and the Leiden algorithm appears to run in *O*(*n* log *n*) on the number of edges of the graph, which is directly related to *k* × *n*. Thus, *k* has an effect both on the performance and the runtime of the method.

To test the actual effects of *k* on the performance of the method, we have applied it to the datasets on Table 1 with varying *k* ∈ {5, 10, 15, 20, 25, 30, 40, 50, 100, 150}. Figure 2 shows an aggregate view of how well the target pathways of each dataset ranked. We see that the effect of *k* on the performance as measure by these results is not very large, we selected *k* = 25 as the methods default value. Unless otherwise specified, we here use *k* = 25 for all our further results.

**Figure 2:**
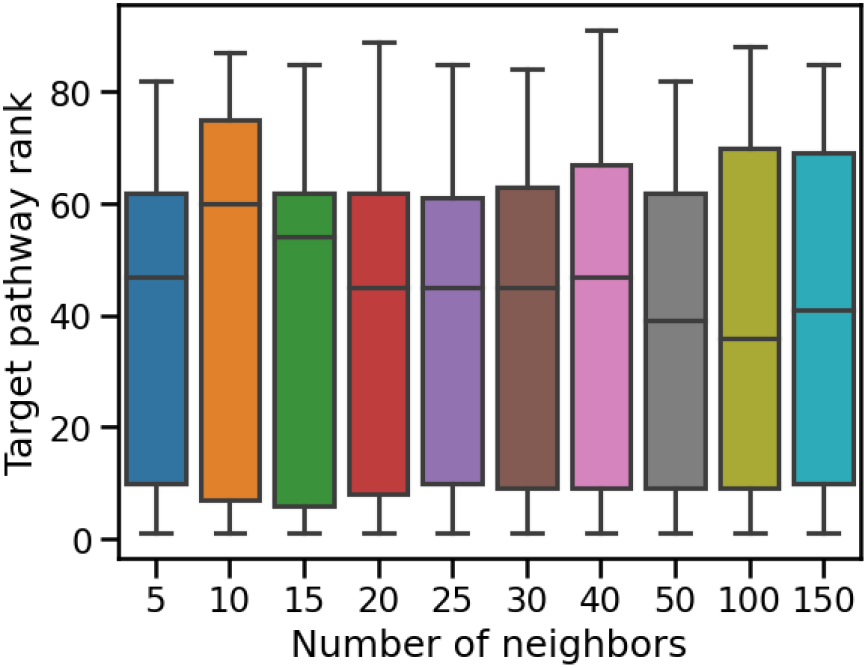
The effect of the selected number of nearest neighbours, *k*, on the methods performance. We plotted the naïve target pathway’s rank performance for different values of *k*. The parameter seems to have little influence over the methods performance so we arbitrary selected *k* = 25.

### MIPath captures a meaningful signal

We want to check if the method can capture a meaningful and stable signal, and one way we can do this is to test if the results are reproducible. To simulate multiple experiments under the same conditions, we can subsample a dataset multiple time, each time performing our method on this reduced dataset, and see how comparable the resulting scores are.

Practically, for every dataset on Table 1 with over 10.000 samples (all but dataset 4), we have subsampled 20% of the dataset 10 times, each time performing our method to score the association between KEGG pathways and the tested phenotype. Having then 10 sets of scores, we calculated the Pearsons’s correlation between every two pairs of sets to get an average correlation over all subsamples. We have observed that the results between runs closely align, with the average pairwise Pearson’s correlation between subsamples of the same dataset being very high at 0.96.

We have also performed permutation tests by permuting the sample labels, confirming our theoretical assumption that by random the expected score is 0.

### MIPath successfully identifies target pathways and benchmarks well against other pathway analysis methods

As a benchmark, we compare our method with both GSEA [1] and Enrichr [42], two widely used methods of pathway analysis. We do this using the datasets on Table 1, their target pathways and tested phenotypes.

For each dataset, we run our method scoring each pathway using the annotation of samples provided by the tested phenotypes and rank the results by their AMI score.

GSEA and Enrichr require a previous statistical analysis to establish differential expression of genes between groups. Since we are allowing more than one group, we perform a one-way ANOVA for each gene and get a significance value for the differential expression of the gene. This ranked list of *p* values is used as an input to GSEA. The method calculates an enrichment score and compares it to 10,000 random permutations to obtain a *p* value for each pathway. The results are then ranked by their *p* value.

Enrichr needs as an input a set of genes, rather than a ranked list, so we need to divide the list into differentially expressed and not differentially expressed genes. For this we first control for multiple testing by converting the *p* values into *q* values following Storey and Tibshirani [43], and threshold the list at a *q* value of 0.05. The set of differentially expressed genes is imputed into Enrichr and its results are ranked by *p* value.

Table 2 shows the results of the benchmark, and we see that our method has the lowest average rank for the target pathways, and outperforms both methods in 4 out of the 9 datasets using this metric. All three methods have a similar standard deviation on the ranks. We note that only one target pathways fell below the 0.05 *p* value threshold with GSEA.

**Table 2:** A comparison of the performance between MIPath, GSEA and Enrichr, based on the rank of a naïvely selected target pathway. An asterisk indicates that the target pathway is did not pass the significance threshold of FDR = 0.05

| Dataset | MIPath | GSEA | Enrichr |
| --- | --- | --- | --- |
| 1 | 85 | 112* | 83 |
| 2 | 11 | 38* | 6 |
| 3 | 61 | 159* | 39 |
| 4 | 45 | 156* | 44 |
| 5 | 48 | 167* | 20 |
| 6 | 10 | 100* | 103 |
| 7 | 1 | 17* | 33 |
| 8 | 5 | 177* | 18 |
| 9 | 84 | 108* | 90 |
| Average | 38.9 | 114.3 | 48.4 |

A more careful examination of the results from dataset 7, which consists of more than 8000 cell from the hematopoietic cells partitioned into 23 cell types [40]. Here the target pathway “Hematopoietic cell lineage” was found to have the largest MIPath score and hence the major differentiating pathway. The second most differentiating pathway was “Cell adhesion molecules (CAMs)”, which indeed should be a differentiating factor between the different cell types in the immune system, e.g. helper T cells and B-cells. The third most important pathway was “Cytokine-cytokine receptor interaction”, which is yet another pathway involved in the immune system, and hence differentiated over the different cell types in the hematopoietic cell linage. This can be contrasted with the top ranked pathways of GSEA, which are “Pantothenate and CoA biosynthesis”, “Asthma”, and “Escherichia coli infection”, which all are less obviously associated with the difference in cell functions of the hematopoietic cell linage.

Figure 3 provides an intuitive illustration of the MIPath scores. We see in Figure 3A that the cell types are clearly separated by the genes in the top scoring pathway, but the same pathway in Figure 3B does not separate well the samples according to which part of the body they were taken. Figures 3C and D show how the separation of cell types look with lower scoring pathways.

**Figure 3:**
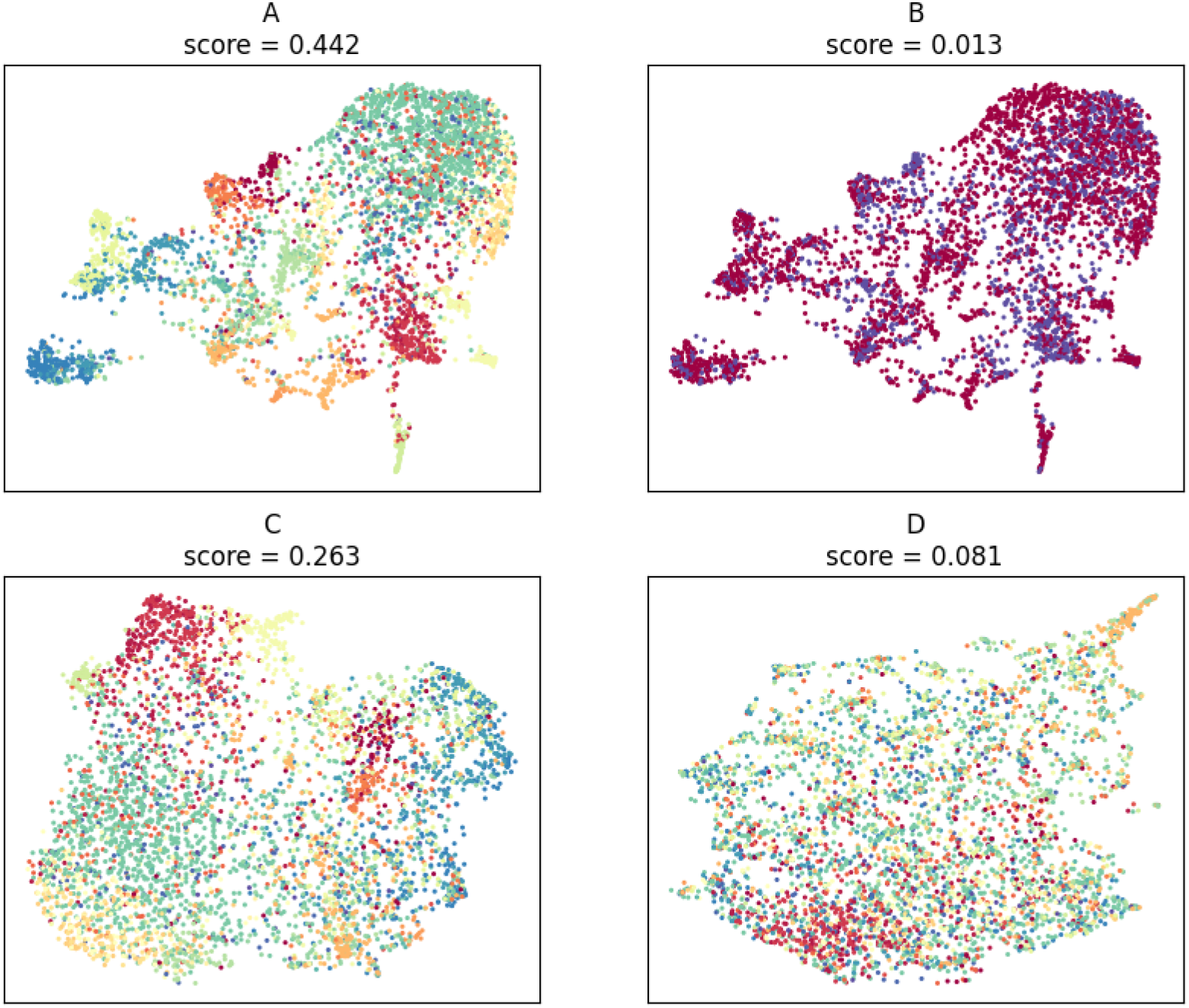
High-scoring pathways indicates a better separation of the samples. We made UMAP plots of the distribution of samples in dataset 7 according to three pathways. The samples were scored and colored for the relation between the “Hematopoietic cell lineage” pathway and A) the inferred cell type and B) the organ the sample were harvested from (bone marrow or liver). The relation between the inferred cell type and the C) “JAK-STAT signaling” pathway and D) the “Lysine degradation” pathway were investigated. Above each plot the MIPath score for the relation is given. The high-scoring relations are visually more heterogeneous than the low-scoring relations.

Turning to dataset 2, which consists of about 17000 cell’s from substantia nigra and cortex of Parkinson’s disease patients. Here we tested against the organism part (substantia nigra vs cortex) and we see that the artificial target Pathway “Parkinson’s disease” is only ranked 11 by MIPath. However, the by MIPath top ranked pathway, “Focal adhesion”, is recently proposed to be associated with Parkinson’s disease [44] and so is the second ranked pathway, “MAPK Signaling” [45].

### The method provides a natural way to deal with confounders

Since most gene expression experiments, especially in single cells, are observational, when studying the association between gene expression and phenotype one must always be aware of the possible presence of confounders. Confounding variables affect both gene expression and phenotype, causing a spurious relationship between the two that may be interpreted as a causal relationship if not properly studied.

Because our method requires only the marginal and joint distributions of the variables to output a score, there is a very natural way to deal with confounders by the use of the respective conditional distributions. By conditioning both marginals and the joint on the possible confounder, we are effectively stratifying the data, as shown by Equation 3, and thus removing the effect of such a confounder from our scores.

One of the most clear examples of confounders that applies to pathway analysis is the patient age in the context of survival analysis. Since the survival variable is measured as the time between measurement and the decease event, age explains a lot of the variance in this variable, but age also affects many biological processes in the cell. Thus, if one intents to study the effects of different pathways on the survival of patients on some disease, they must be aware of the spurious relationship between those two that might be caused by the age of the patients.

We tested how our model can handle this case on a breast cancer dataset. For this, we downloaded gene expression data from the METABRIC dataset, consisting of microarray reads from 1992 breast cancer specimens, together with the clinical annotation containing the survival information and age of the respective patients. We then classify the patients into 5 quantiles for both age and survival time, and after standardizing the gene expression, we run our method to test the association between all Reactome pathways and the survival quantiles, as well as the same association when conditioning on the age quantiles.

If we rank the pathways with the biggest difference between the unconditioned score and the one where we use the conditional distributions on age, we can find the ones where the spurious relationship caused by age is strongest. Among the top 1% of pathways with the largest drop in score are: Inhibition of nitric oxide production [46], Defective Mismatch Repair Associated With MSH6 [47], Signaling by Erythropoietin [48], ABC transporter disorders [49], Biosynthesis of protectins [50] and Elevation of cytosolic Ca2+ levels [51], all of which have been shown to be associated with aging. Other well known pathways related to aging, Extension of Telomeres [52] and Serotonin and melatonin biosynthesis [53], are also among the top 3%.

## Conclusion

In this paper, we have presented a new method for pathway analysis, MIPath, that differs from others for its use of concepts in information theory to score the association between pathway activity and phenotype. Information theory as a field has long dealt with the problem of analysing signals in complex systems, and as such is very suitable for applications in molecular biology [54]. Practically, we find modules of samples for each pathway and score the association between these modules and any phenotype classification by using a mutual information score that has been normalized and corrected for randomness.

This process has several advantages. First, it does not rely on any kind of previous statistical analysis to rank or select significant genes, as it identifies sample modules directly from the gene expression. By using the subspace of gene expression defined by each pathway, and converting it to a nearest neighbor graph (with shared neighbors weights), we avoid performing the module detection step on spaces of different dimensionality. This is important as many clustering methods are known to have varying performance depending on the dimensionality of the space [17].

Second, by scoring using mutual information, and embedding the hypergeometric error model in the calculation of each individual score, we get a very sensitive model that can detect even weak signals at high significance levels. This, combined with the score normalization, also make the scores comparable across experiments. An added benefit is the possibility of using conditional mutual information to deal with confounding factors.

Third, it has only one parameter, namely the number of neighbors used for the graph construction, which had an intuitive meaning as the tradeoff between global and local structure of the samples. Practically, however, it does not have a big effect on the performance of the method, thanks in part to the use of the Shared Nearest Neighbors similarity metric.

Lastly, this test is self-contained, meaning that the scores for each pathway depends only on the expression of genes belonging to that pathway, and does not rely on a gene expression background, which further increases its sensitivity [55].

We have shown that this method captures a stable signal, and that it perform well by measuring how highly known targeted pathways are ranked in nine different datasets, and comparing it to the ranks obtained by both GSEA and Enrichr under the same conditions. We have also shown that conditional mutual information is can successfully deal with confounders.

There are, however, some drawbacks for the model. The biggest being that it does not handle continuous variables, and variables such as survival or age have to be discretized into quantiles in order to be scored. This stems from the fact that we rely on empirical distributions to be able to calculate the mutual information. Even the gene expression itself is also discretized in a way, with samples that are close together in a pathway subspace considered equal, which results in a theoretical loss of information.

Another important limitation is that due to the use of the Leiden algorithm to find sample modules, our method requires more samples than others to produce a meaningful result. Further improvements could be made here, with alternate methods for finding the sample modules depending on the set size, or even allowing for more omics modalities to be included.

## Code availability

The source code for MIPath can be found at https://github.com/statisticalbiotechnology/mipath

## Acknowlegments

The computations were enabled by resources provided by the Swedish National Infrastructure for Computing (SNIC) at HPC2N.

