## Supplemental Information for "Pathway Analysis Through Mutual Information"

[1]Science for Life Laboratory, KTH Royal Institute of Technology
[*]Corresponding

#### Expected conditional mutual information

Nguyen et al.[1] shows us how to calculate the expected mutual information over two random partitions of the data using a hypergeometric model for randomness. To extend this result to the conditional mutual information, we start with the definition of $I(X, Y|Z)$ for the discrete variables $X$, $Y$ and $Z$ with support $\mathcal{X}$, $\mathcal{Y}$ and $\mathcal{Z}$ respectively:

$$I(X, Y|Z) = \sum_{z\in\mathcal{Z}} p_Z(z) \sum_{x\in\mathcal{X}} \sum_{y\in\mathcal{Y}} p_{X,Y|Z}(x, y|z) \log\left(\frac{p_{X,Y|Z}(x, y|z)}{p_{X|Z}(x|z)p_{Y|Z}(y|z)}\right) \quad (1)$$

and apply the same convention to its joint contingency table. We now have the tensor $M = [n_{ijk}]$ where $i = 1...R$, $j = 1...C$ and $k = 1...S$, $A = [a_{ik}]$ and $B = [b_{jk}]$ are the 2-way tables of $(X, Z)$ and $(Y, Z)$ respectively and $c = (c_1, ..., c_k)$ is the marginal of $Z$. We can then rewrite Eq. 1 as:

$$I(M) = \sum_{k=1}^{S} \frac{c_k}{N} \sum_{i=1}^{R} \sum_{j=1}^{C} \frac{n_{ijk}}{c_k} \log\left(\frac{c_k n_{ijk}}{a_{ik} b_{jk}}\right)$$

Where $N$ is the sum of all elements in the table. We want to calculate its expectation over the set $\mathcal{M}$ of all possible tensors $M$ that result in the same $A$ and $B$

$$E\{I(M)|A, B\} = \sum_{M\in\mathcal{M}} \sum_{k=1}^{S} \frac{c_k}{N} \sum_{i=1}^{R} \sum_{j=1}^{C} \frac{n_{ijk}}{c_k} \log\left(\frac{c_k n_{ijk}}{a_{ik} b_{jk}}\right) P(M|A, B)$$

Here, we note that in our model of randomness, $c$ is fixed, so we can rearrange following Nguyen et al.:

$$E\{I(M)|A, B\} = \sum_{k=1}^{S} \frac{c_k}{N} \sum_{i=1}^{R} \sum_{j=1}^{C} \sum_{n_{ijk}} \frac{n_{ijk}}{c_k} \log\left(\frac{c_k n_{ijk}}{a_{ik} b_{jk}}\right) P(M|n_{ijk}, A, B)$$

We also note that due to the lack of constrains on the $(X, Y)$ contingency table, $P(M|n_{ijk}, A, B)$ only depends on elements $a_{ik}$ and $b_{jk}$ of $A$ and $B$. Thus we have.

$$E\{I(M)|A, B\} = \sum_{k=1}^{S} \frac{c_k}{N} \sum_{i=1}^{R} \sum_{j=1}^{C} \sum_{n_{ijk}} \frac{n_{ijk}}{c_k} \log\left(\frac{c_k n_{ijk}}{a_{ik} b_{jk}}\right) P(M|n_{ijk}, a_{ik}, b_{jk})$$

And $P(M|n_{ijk}, a_{ik}, b_{jk})$ can be calculated [1] as:

$$P(M|n_{ijk}, a_{ik}, b_{jk}) = \frac{\binom{c_k}{n_{ijk}}\binom{c_k - n_{ijk}}{a_{ik} - n_{ijk}}\binom{c_k - a_{ik}}{b_{jk} - n_{ijk}}}{\binom{c_k}{a_{ik}}\binom{c_k}{b_{jk}}}$$

Adding that $n_{ijk}$ can only take values between $\min(0, a_{ik} + bjk - c_k)$ and $\min(a_{ik}, b_{jk})$ we have the final expression for the expectation

$$E\{I(M|A,B)\} = \sum_{k=1}^{S} \frac{c_k}{N} \sum_{i=1}^{R} \sum_{j=1}^{C} \sum_{n_{ijk}=\min(0, a_{ik}+bjk-c_k)}^{\min(a_{ik}, b_{jk})} \frac{n_{ijk}}{c_k} \log\left(\frac{c_k n_{ijk}}{a_{ik} b_{jk}}\right) \frac{a_{ik}! b_{jk}! (c_k - a_{ik})! (c_k - b_{jk})!}{c_k! n_{ijk}! (a_{ik} - n_{ijk})! (b_{jk} - n_{ijk})! (c_k - a_{ik} - b_{jk} + n_{ijk})!}$$

### References

[1] Xuan Vinh Nguyen, Julien Epps, and James Bailey. Information theoretic measures for clusterings comparison: is a correction for chance necessary? In *ICML*, 2009.
